# Identification of the MRTFA/SRF pathway as a critical regulator of quiescence in cancer

**DOI:** 10.1101/2024.11.15.623825

**Authors:** Santiago Panesso-Gómez, Alexander J. Cole, Alyssa Wield, Vivian I. Anyaeche, Jaynish Shah, Qi Jiang, Tonge Ebai, Allison C. Sharrow, George Tseng, Euisik Yoon, Daniel D. Brown, Amanda M. Clark, Scott D Larsen, Ian Eder, David Gau, Partha Roy, Kris N. Dahl, Lam Tran, Hui Jiang, Priscilla F McAuliffe, Adrian V Lee, Ronald J. Buckanovich

## Abstract

Chemoresistance is a major driver of cancer deaths. One understudied mechanism of chemoresistance is quiescence. We used single cell culture to identify, retrieve, and RNA-Seq profile primary quiescent ovarian cancer cells (qOvCa). We found that many qOvCa differentially expressed genes are transcriptional targets of the Myocardin Related Transcription Factor/Serum Response Factor (MRTF/SRF) pathway. We also found that genetic disruption of MRTF-SRF interaction, or an MRTF/SRF inhibitor (CCG257081) impact qOvCa gene expression and induce a quiescent state in cancer cells. Suggesting a broad role for this pathway in quiescence, CCG257081 treatment induced quiescence in breast, lung, colon, pancreatic and ovarian cancer cells. Furthermore, CCG081 (i) maintained a quiescent state in patient derived breast cancer organoids and, (ii) induced tumor growth arrest in ovarian cancer xenografts. Together, these data suggest that MRTF/SRF pathway is a critical regulator of quiescence in cancer and a possible therapeutic target.

**Significance:** Quiescence is a critical driver of chemoresistance. The MRFT-SRF pathway regulates cancer cell quiescence and inhibiting the MRTF-SRF pathway can prevent the outgrowth of quiescent cancer cells and improve cancer outcomes.

## Introduction

While targeted therapeutics and immunotherapy have had a significant impact in many tumor types, for many patients, chemotherapy remains the mainstay of treatment. Unfortunately, for many cancers such as lung and ovarian, despite an initial response to chemotherapy, relapse is common and typically occurs within 2 years of completing therapy ^1^. This suggests an inherent mechanism of chemoresistance. A better understanding of mechanisms of chemotherapy resistance is critical to inform the development of therapies and to improve patient outcomes.

One poorly studied mechanism of chemotherapy resistance is quiescence. Quiescent cells reversibly exit the cell cycle and are thus refractory to standard chemotherapies which preferentially target rapidly proliferating cells ^2^. Quiescent cancer cells, often a subset of cancer stem-like cells (CSCs), exhibit resilience to chemotherapy and other stressors ^3,4^: slow cycling pancreatic CSCs demonstrate enhanced chemotherapeutic resistance and tumor initiation capacity ^5^, quiescent leukemia stem cells ^6,7^, SOX2+ medulloblastoma stem cells ^8^, and colon cancer stem cells ^9–11^ have all been associated with therapeutic resistance and tumor recurrence.

Ovarian cancer is a disease in which chemotherapy resistance is particularly problematic. Chemotherapy remains the standard adjuvant therapy for patients with ovarian cancer and despite an almost complete clinical response with surgery and chemotherapy, approximately 60-70% will relapse and ultimately succumb to chemo-resistant disease ^12^. Quiescence is emerging as critical driver of chemotherapy resistance in ovarian cancer. Gao et. al. identified a population of slow cycling, chemotherapy-resistant OvCa CSCs that drive cancer recurrence in primary human tumors ^13^. Similarly, studies with OvCa patient samples and PDX models indicate that residual OvCa cells are enriched for Ki67(-) CSCs ^14,15^. Furthermore, when OvCa cells are grown in suspension, commonly seen with OvCa associated ascites, they assume a reversible quiescent stem-like state and are chemotherapy resistant ^16^.

We recently demonstrated that the transcription factor Nuclear Factor of Activated T-cells-C4 (NFATC4) drives OvCa cell quiescence and chemotherapy resistance ^17^. This is analogous to the role of NFATc1 in promoting hair follicle stem cell quiescence ^18–20^. In addition, we found that inherently resistant quiescent ovaria cancer cells (qOvCa) secrete factors, such as follistatin, to increase chemotherapy resistance in neighboring non-quiescent cells ^21^. Validating quiescence as a therapeutic target in ovarian cancer, we found that loss of follistatin activity could improve response to chemotherapy and increase cure rates in animal models ^21^.

To further increase our understanding of drivers of quiescence and identify putative quiescent cell therapeutic targets/pathways, here, we used single-cell microfluidic culture of primary ovarian cancer cells to identify and perform single-cell RNA sequencing on qOvCa. We identified numerous proteostasis related pathways impacted in qOvCa and found that the Myocardin-Related Transcription Factor-A (MRTFA)/Serum Response Factor (SRF) pathway is a master regulator of many qOvCa expressed genes. Inhibition of MRTFA/SRF signaling with the small molecule CCG257081 induced a quiescent state in a range of cancer types including ovarian, breast, lung, colon, and pancreatic cancers. Profiling of CCG257081 treated cells revealed additional therapeutic pathways (such as polio-like-kinase and aurora kinase) which, when targeted did not induce cell death, but rather induced quiescence. Finally, treatment of ovarian cancer xenografts with a maintenance dose of CCG257081 forced quiescence in cancer cells, significantly delaying disease progression, and suggesting that inhibition of the MRTFA pathway may function as a novel quiescence maintenance therapy to control recurrent disease by forcing quiescence in residual chemoresistant cells.

## Results

### RNA signature of quiescent ovarian cancer stem-like cells

To identify genes differentially expressed in proliferative vs. non-proliferative phenotype, we used previously described microfluidic single cell culture device with laser retrieval capacity ^22^. FACs isolated ALDH^+^CD133^+^ ovarian cancer cells were loaded into a 10,000 microwell device using gravity flow, and each microwell was imaged to ensure capture of a single cell (**Fig 1A-B**,). Microwells were imaged daily to monitor division, and after 5 days in culture cells were stained with a viability dye. While ∼90-95% of cells underwent at least one cellular division, with most undergoing numerous divisions during the 5-day period, ∼5% of the viable cells did not proliferate, remaining as single cells (**Fig 1B**). We used laser retrieval to isolate ∼200 dividing cells and ∼100 quiescent cells from two primary ovarian cancer patient cell lines PT340 and PT412. Isolated cells were then profiled using the CellSeq2 single cell RNA-sequencing method ^23^.

**Figure 1.**
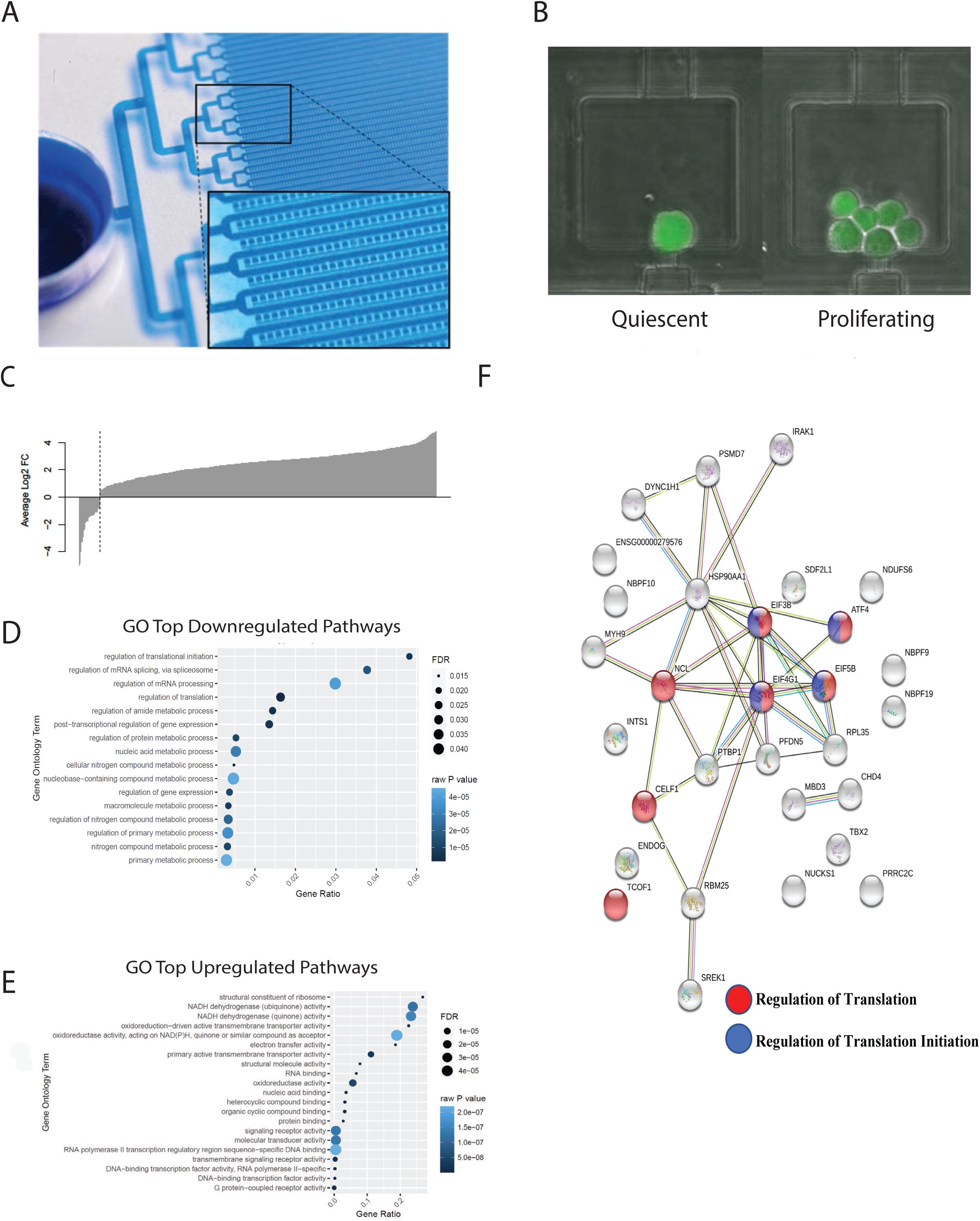
Identifying and single-cell-RNA-Seq profiling quiescent ovarian cancer stem cells identifies key quiescence genes and pathways. **A.** Image displaying the single-cell microfluidic culture platform with photo-mechanical cell detachment and retrieval capacity used to identify and isolate quiescent ovarian cancer cells. **B.** Images of live cells after 5 days in culture, identifying (left) a single non-dividing/quiescent cell and (right) a well with multiple cells that derived from a single proliferating cell. **C.** Waterfall plot of differentially expressed gene in quiescent cells vs proliferating cells. **D** and **E.** GO analysis of the top down- and upregulated molecular pathways. **F.** STRING analysis to the significantly enriched downregulated genes.

Despite the limited depth of sequencing using the manual CellSeq2 amplification approach, we identified hundreds of genes which were statistically significant differentially expressed between the proliferating and quiescent cells (**Fig 1C**, **Table 1)**. Gene-Ontogeny (GO) analysis of downregulated genes demonstrated significant downregulation of pathways associated with cellular metabolism, protein translation, RNA processing and metabolism, and chromatin binding (**Fig 1D),** while the major up regulated pathways were NADH/NADPH activity, transmembrane transport, and RNA binding (**Fig 1E).** Due to the abundance of upregulated genes, we decided to investigate the function of the more modest number of significantly downregulated genes. STRING (Search Tool for the Retrieval of Interacting Genes/Protein) analysis of the downregulated genes identified EIF4G1, MHY9 and NCL as key signaling factors (**Fig 1F)**^24^.

**Table 1.** Top downregulated targets in Sc-RNAseq of proliferating vs. non-proliferating cells.

| Protein Translation | RNA Regulation | Chromatin Remodeling and DNA Damage Response |
| --- | --- | --- |
| EIF5B (2.8)** | TCOF1 (6.3)* | MALAT1 (13.4)** |
| EIF3B (2.0)* | SREK1 (6.2)* | MBD3 (9.4)** |
| RPL35 (2.0)* | RBM25 (6.1)* | TCOF1 (6.3)* |
| EIF4G1 (2.0)* | ENDOG (5.3)* | NUCKS1 (3.3)* |
|  | INTS1 (3.5)* | CHD4 (3.1)** |
|  | NCL1 (2.8)** |  |
|  | CELF (2.5)* |  |
|  | PTBP1 (2.0)* |  |

### Downregulation of regulators of translation induces a quiescent state

To determine whether downregulation of EIF4G1, MHY9 or NCL promotea a quiescent phenotype, we reduced their expression using siRNA (**Sup Fig 1A-B**) in the PT340 and PT412 OvCa cell lines, ^21^. Compared to scrambled siRNA control, siRNA downregulation of all target genes resulted in a significant decrease in cell number without impacting cell viability (**Fig 2A, Sup Fig 1C-D,** p<0.05). Cell cycle analysis showed that for all three genes, siRNA downregulation resulted in a significant increase in cells in the G0/G1 phase of cell cycle (**Fig 2B**, p<0.01). This effect was corroborated using the FUCCI cell cycle reporters, which demonstrated that downregulation of each gene significantly increased p27KIP1, which has been strongly linked with quiescence (**Fig 2C-D**, p<0.01). MYH9 and NCL knockdown, but not EIF4G1 knockdown, increased the p27/CDT1+ cells felt to be in G0 (**Fig 2E**, p<0.01).

**Figure 2.**
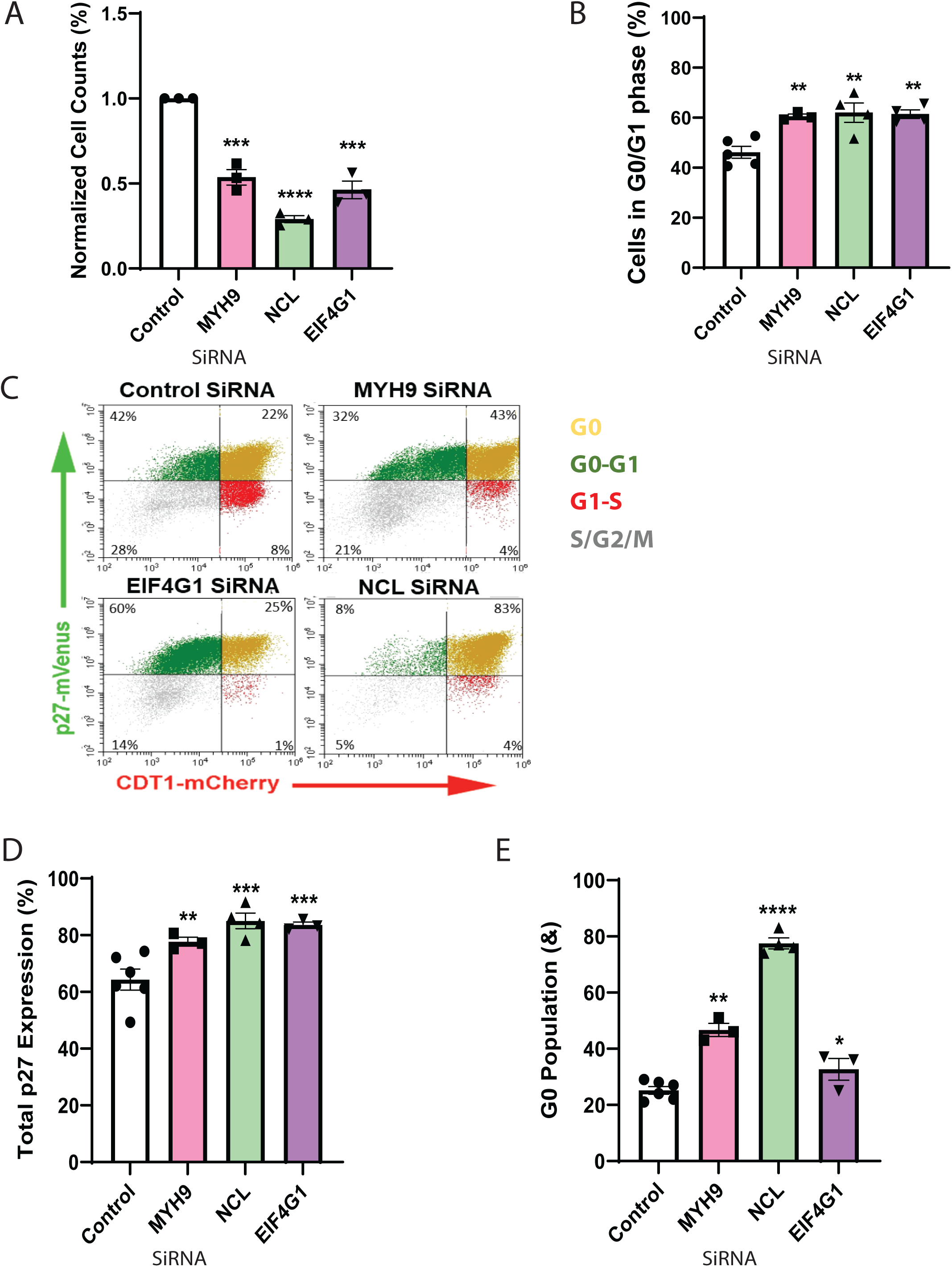
Downregulation of EIF4G1, MHY9, and NCL contributes to quiescence. (**A)** Cell counts and **(B)** cell cycle summary of PT412 cells treated with EIF4G1, MHY9, or NCL siRNA compared to a scrambled control. **C.** FUCCI cell cycle analysis of HEY1 cells treated with scrambled control siRNA or EIF4G1, MHY9, or NCL siRNA. **D.** Quantification of total p27 expression of SiRNA knockdown compared to control. **E.** p27KIP1-mVenus and CDT1-mCherry-positive cells in G0, showing an increase in SiRNA knockdowns as compared to control. Each experiment was repeated, in triplicate, a minimum of three times. Results were compared with Student’s T-test, *p<0.05, **p<0.01.

### Therapeutic benefits of targeting translation

As both MHY9 and NCL downregulation resulted in the induction of cell growth arrest and an apparent G0 state we next tested the impact of MHY9 and NCL inhibitors, blebbistatin and oridonin respectively, on ovarian cancer cells ^25^. Both drugs significantly reduced cell number in a dose dependent manner (**Fig 3Ai and 3Bi,** p<0.001), without affecting cell viability (**Fig 3Aii and 3Bii,** p<0.0001). To determine whether these drugs, by inducing quiescence, could delay cancer growth in vivo, we tested them in a PT340 xenograft model. Both drugs delayed tumor growth, with NCL inhibition with oridonin having the greatest effect (**Fig 3C-D**. p<0.0001). Immunohistochemical staining for Ki67, in control and treated tumors, confirmed a significant reduction in Ki67+ cells in both treatment groups (blebbistatin, p<0.01 and oridonin, p<0.001), with percent reduction in Ki67 staining mirroring percent reduction in tumor growth (**Fig 3E-F)**. Notably, consistent with a quiescent phenotype, oridonin-treated tumor cells also appeared much smaller. Together, these data suggest that inhibition of either MHY9 or NCL induce a quiescent phenotype that can restrict tumor growth.

**Figure 3.**
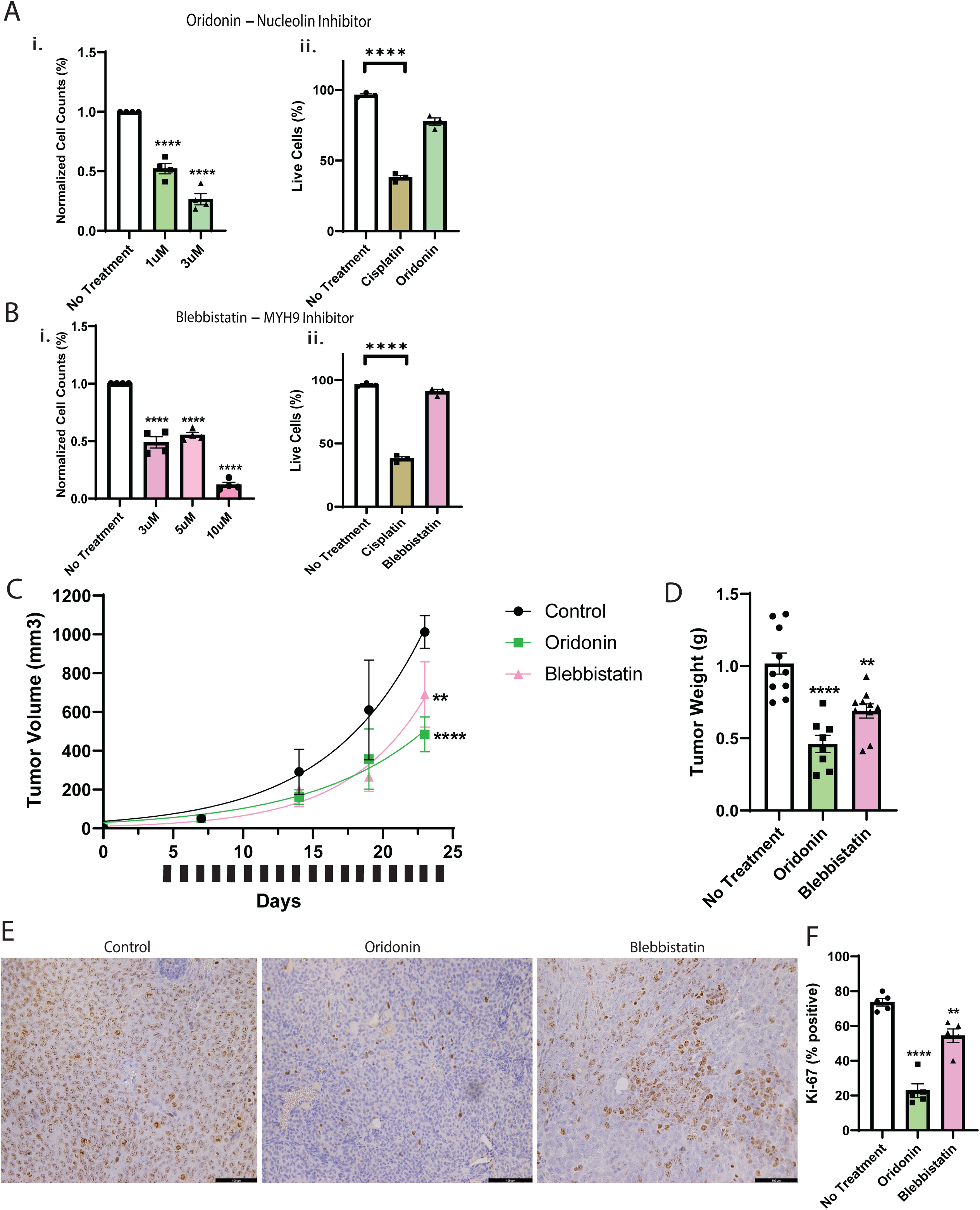
Chemical inhibition of MHY9 and NCL reduce cell proliferation *in vitro* and *in vivo*. **A-B.** Normalized (i) cell counts and (ii) viability percentage in PT340 cells line treated with the indicated doses of oridonin or blebbistatin for 72 hours. Three µM cisplatin was used as a cell death viability control. **C-D.** Tumor growth curve and final tumor weights of PT340 tumor xenografts (n=10/treatment group). Black bars indicate treatment with 20mg/Kg oridonin, 2mg/Kg blebbistatin, or DMSO control daily. **E-F.** Representative images and quantification of Ki67 staining. *In vitro* assays were replicated at least three times. Tumor Ki67 was analyzed for 6 to 8 independent sections of 3 independent tumors for each treatment group, *p<0.05, **p<0.01, ***p<0.001, ****p<0.0001.

### Linking the SRF/MRTF pathway and quiescence

While drugs targeting both MYH9 and NCL proved effective in reducing tumor growth, neither had a profound effect. We speculated that resistance is more likely when targeting a single downstream mediator of a quiescent phenotype and therefore sought to identify a more global regulator of quiescence. By mining the harmonizome genes and protein database, we identified that numerous genes, including EIF4G1, MHY9 and NCL, as being transcriptional targets of the MRTF/SRF pathway (https://maayanlab.cloud/Harmonizome/gene_set/SRF/ENCODE+Transcription+Factor+Targets). Indeed, 61% of qOvCa differentially expressed genes are ChIP-seq validated SRF targets (**Fig 4A**) ^26–30^, suggesting that this pathway is a global regulator of quiescence/proliferation and could therefore be a better therapeutic target.

**Figure 4.**
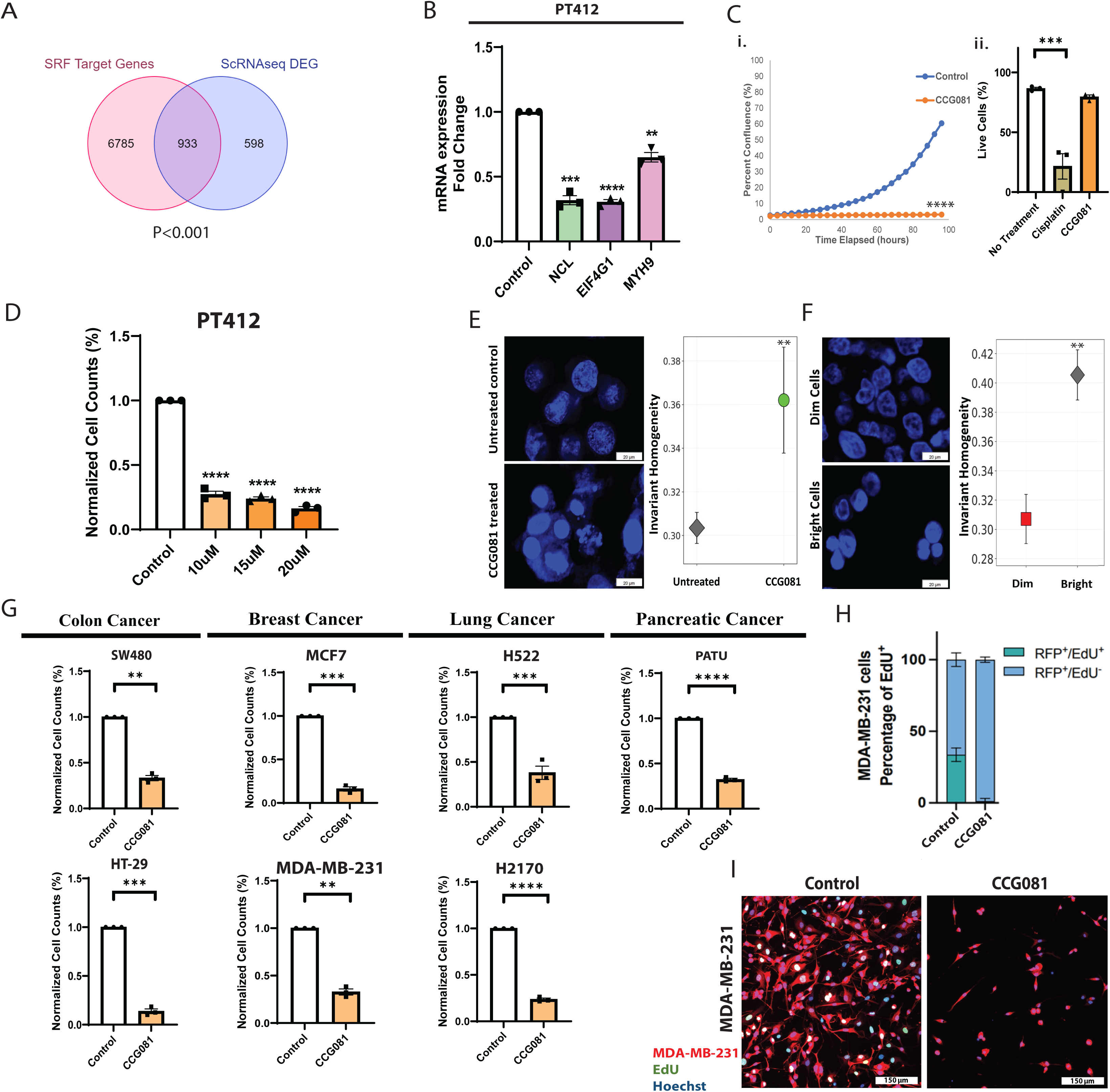
The SRF/MRTF transcriptional inhibitor CCG257081 acts through NCL, MHY9 and EIF4G1 to induces a quiescent state in cancer cells. **A.** Venn diagram showing overlap of differentially expressed genes of scRNAseq of primary quiescent cells and previously reported SRF targets ^26^. **B**. Relative mRNA expression for the indicated genes as assesses by qRT-PCR in PT412 cells treated with DMSO (control) or CCG257081 for 72-h. **C**. IncuCyte real time imaging analysis of cellular confluence and viability analysis of control and CCG257081 (15µM every 48 h for 4 days). **D.** Normalized cell counts of PT412 treated with CCG257081 or Vehicle control (DMSO). **E-F.** DAPI chromatin IF and measure of nuclear invariant homogeneity for PT340 cell line (E) treated with DMSO or CCG257081 or, (F) isolated CFSE labeled slow/non-dividing (Bright) or rapidly proliferating (Dim) cells. **G.** Normalized cell counts of colon (SW480, HT-29), breast (MCF7, MDA-MB-231), lung (H522, H2170), and pancreatic (PATU) cell lines, treated with DMSO or CCG257081. **H-I.** Summary and representative IF image of EdU staining in control and CCG257081-treated RFP-MDA-MB231. Except were otherwise indicated, CCG257081 treatment was 15µM every 48 hours for 72 hours. *In vitro* assays were replicated in triplicate at least three times and compared with an ANOVA, *p<0.05, **p<0.01, ***p<0.001, ****p<0.0001.

To determine if the MRTF/SRF axis regulates the genes associated with quiescence, we treated the ovarian cancer cell lines PT340, PT412 and OVSAHO with CCG257081, a high-affinity small-molecule inhibitor of SRF/MRTF transcriptional activity ^31,32^. qRT-PCR of treated cells indicated CCG257081 significantly repressed mRNA expression of EIF4G1, MHY9 and NCL (**Fig 4B, Supp. Fig 2A,** P<0.05).

We next tested the impact of CCG257081 treatment on proliferation/quiescence. Consistent with the induction of quiescence, real time live cell imaging using the Incyte revealed CCG257081 treatment restricted cell proliferation (**Fig 4Ci-D**), which was confirmed using cell counts while having no effect on cellular viability (**Fig 4Cii**, **Supp. Fig 2C**).

The chromatin of quiescent cells is generally tightly compacted ^33^. To confirm a similar phenotype in qOvCa cells we isolated and imaged vital dye retaining (bright) quiescent cells and vital dye diluting (dim) proliferating cells. In parallel, we imaged CCG257081 treated and control cells. Both the “bright” cells and the CCG257081 treated cells demonstrated a similar and significant increase in chromatin invariant homogeneity, suggestive of chromatin condensation (**Fig 4E-F**).

To determine whether the impact of CCG257081 on cell proliferation was ovarian cancer specific or applicable across different tumor types, we treated numerous cancer types with CCG257081, including colon (SW480, HT-29), breast (MCF7, MDA, T47D), lung (H522, H2170), and pancreatic (PATU). We found that CCG257081 restricted cell growth in all cancer cell lines without impacting cell viability (**Fig 4G, Supp. Fig 2D**), suggesting induction of a quiescent state. Further supporting the induction of quiescence, treatment of RFP labeled MDA-MB-231 cells (triple-negative breast cancer cell line) with CCG257081 in the presence of EdU to assess proliferation demonstrated that CCG257081 nearly eliminated for EdU uptake (**Fig 4H-I**).

### CCG257081 acts through p27 to induce quiescence

We next evaluated the impact of CCG257081 on the cell cycle using the G0/G1 p27KIP1- mVenus/CDT1-mCherry Fucci reporter vectors ^34^. We observed a significant increase in cells expressing the p27-mVenus (**Fig 5A-B**). To determine whether endogenous p27 levels are increased, we performed qRT-PCR and western blotting on CCG257081-treated cells. We observed an increase expression at the mRNA level (**Fig 5C**), as well as a dose-dependent increase in endogenous p27 protein levels with CCG257081 treatment (**Fig 5C-E**). Supporting the FUCCI data, using propidium iodide (PI) staining we corroborated an increase in the G0/G1 population in CCG257081 treated cells (**Supp. Fig 3A**).

**Figure 5.**
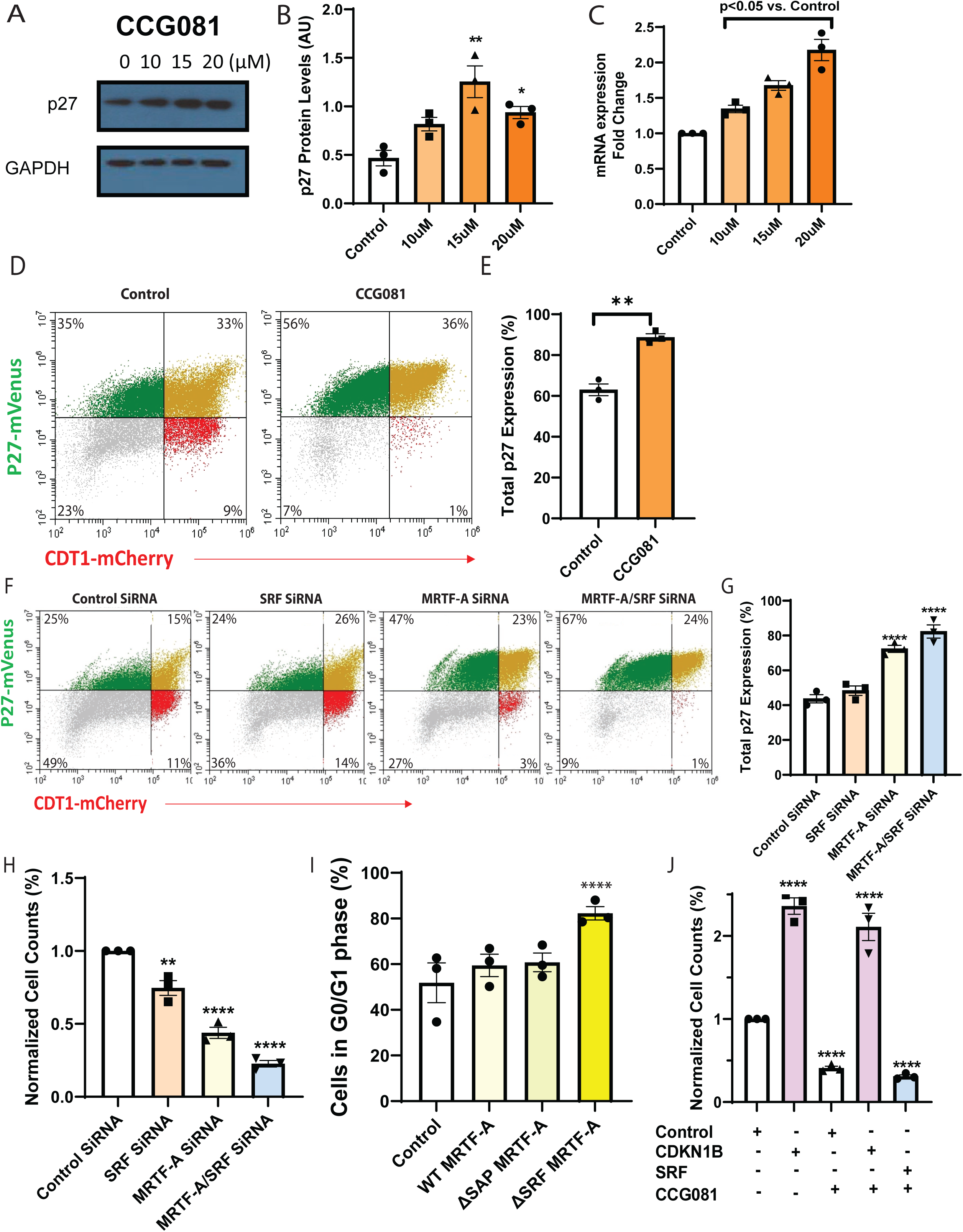
CCG257081 acts through p27 to induce quiescence. **A-B** p27 (p27kip1) Western blot and densitometry in control and CCG081-treated cells. **C**. p27 Relative mRNA expression in control and CCG081-treated PT340 cells. **D-E.** FUCCI cell cycle analysis of control and CCG081- treated HEY1 cells. **F-G**. FUCCI cell cycle analysis and quantification of p27-mVenus expression in HEY1 cells treated with scrambled control, MRTFA, SRF, or MRTFA+ SRF siRNA. **H.** Cell counts of PT340 cells treated with the indicated SiRNAs. **I.** Percent of cells in G0/G1 as measured by PI staining in Pt340 cells transfected with wild type (WT) MRTF-A, MRTF-A with a deletion of the SAP domain (ΔSAP), or MRTF-A with a deletion of the SRF binding domain (ΔSRF). **J.** Cell counts of PT340 cells treated with the indicated compounds alone or in combination with 15 µM of CCG257081. In all experiments, CCG257081 treatment was every 48 hours for 3 days. All assays were replicated, in triplicate, at least three times and compared with an ANOVA, *p<0.05, **p<0.01, ***p<0.001, ****p<0.0001.

To link the CCG081 effect to the MRTFA/SRF pathway, we used siRNA knockdown. We observed that MRTF-A knockdown significantly increased the number of p27+ cells and cells in G0. Combination knockdown of MRTF-A and SRF further increased the number of p27+ cells/cells in G0, closely mirroring the effect of CCG257081 treatment (**Fig 5F-H, Supp. Fig 4A)**. Next, we used previously established MDA-MB231 cells expressing either wild type MRTF-A, MRTF-A with a deleted SRF binding domain (MRTF-ΔSRF), or MRTF-A with a deleted SAP DNA binding domain (MRTF-ΔSAP) and performed PI based cell cycle analysis. Only cells expressing the MRTF-ΔSRF demonstrated a significant increase in the number of cells in G0/G1(**Fig 5I).**

To determine whether p27 was necessary for the growth restricting effect of CCG257081, we knocked down CDKN1B (gene which encodes p27) expression with siRNA and subsequently treated with CCG257081. In parallel, we knocked down SRF. We observed that p27 knockdown protected the cells against CCG257081-mediated quiescence confirming the role of p27 (**Fig 5J**, p<0.01, **Supp. Fig 4B**). In contrast, SRF knockdown enhanced CCG257081’s effects. Combined these data indicate that inhibition of the SRF/MRTF pathway, via induction of p27, results in a quiescent state in cancer cells.

### Identifying other quiescent cancer cell therapeutic targets

To identify additional genes being downregulated by CCG257081, we performed RNA sequencing on CCG257081 treated cells from three independent ovarian cancer cell lines. Principle component analysis demonstrated a clear treatment effect (**Supp. Fig 5**). Volcano plot using a Log2 fold change of at least one and an adjusted p value <0.05 revealed that, like the single-cell quiescence signature, more genes were upregulated (994) than downregulated (352) across the three cell lines (**Fig 6A**).

**Figure 6.**
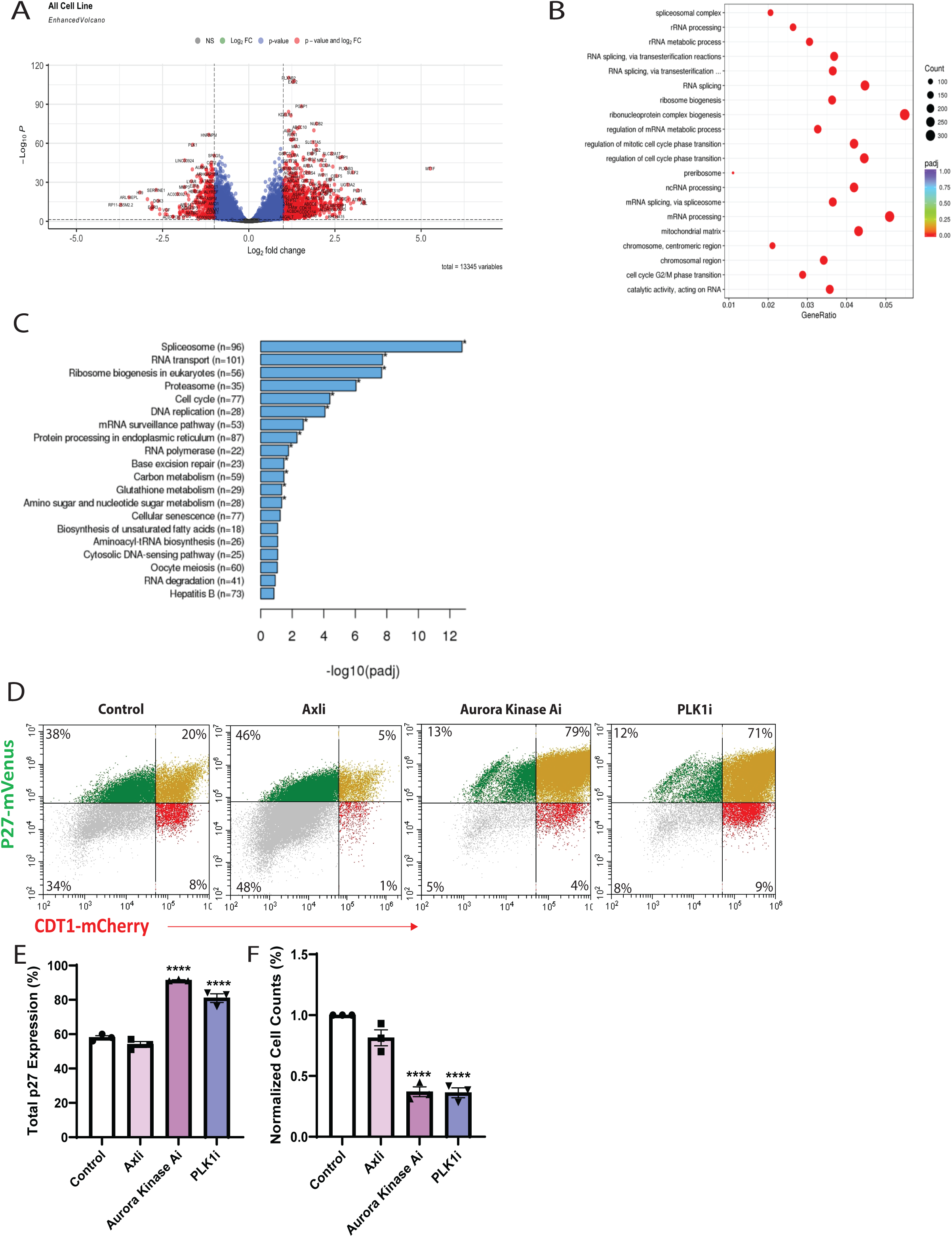
RNA sequencing of CCG257081-treated cells supports induction of quiescence. **A.** Volcano plots of differential gene expression between control and CCG257081-treated cells. **B.** Gene Ontology (GO) analysis of the top involved pathways following CCG257081 treatment. **C.** Kyoto Encyclopedia of Genes and Genomes (KEGG) in involved in CCG257081-treated cells. **D.** FUCCI cell cycle analysis of HEY1 cells treated with either AXL (bemcetinib), Aurora Kinase-A (AURKA, alisertib), or polio-like kinase 1 (PLK1, onvansertib) inhibitors. **E-G.** Percent p27+ cells, normalized cell counts, and percent live cells in cells treated with the indicated inhibitors.

Downregulated genes included numerous cell cycle and DNA damage response linked factors and growth factors including: PLK1, CCNA2, CDC25A, E2F8, AURKA, EGR3, and HB-EGF. Gene ontology pathway analysis of downregulated genes identified changes in pathways related to telomere binding, mitotic and meiotic DNA replication, and DNA repair. Upregulated genes include AXL, SEMA3B, TGFb1L1, and FBXO2. Gene ontology pathway analysis of upregulated genes identified changes in pathways related to peptide catabolism, cell adhesion, lipid and fatty acid metabolism, and several biosynthetic processes (**Fig 6B**). KEGG analysis also showed involvement in various cell functions including the spliceosome, RNA transport, DNA replication and cell cycle (**Fig 6C**)

Several of the genes downregulated in our dataset, such as PLK1 (polio-like kinase 1) and AURKA (aurora kinase-A) have been proposed as cancer therapeutic targets ^35–37^. To determine if inhibition of these targets restricts cancer growth by inducing quiescence (as opposed to inducing cell death), we treated ovarian cancer cells with PLK1 (onvasertib) and AURKA (alisertib) inhibitors. Consistent with induction of quiescence, both inhibitors reduced cell numbers without inducing significant cell death. In addition, Fucci reporter analysis indicated a strong upregulation of p27KIP1+/CDT1+ cells in G0 with the strongest effect observed with the aurora kinase inhibitor (**Fig 6D-E**, p<0.01). Confirming the effect, with the use of both of inhibitors, we also saw a reduction in cell numbers, without affecting their viability (**Fig 6F**, p<0.01, **Supp. Fig 5A**).

### Pharmacologically enforced quiescence as a therapeutic strategy

To evaluate how our compound would translate to a clinical setting, we first treated two triple-negative breast cancer (TNBC) organoids with CCG257081. Treatment resulted in (i) a reduced number, and (ii) size of organoids (**Fig 7A-C**, p<0.05), and (iii) organoids with a significantly lower percentage of Ki67 positive cells (**Fig 7B-D**, p<0.05). Linking this to the MRTF/SRF pathway, we had previously noted that tumors initiation by cells MDA-MB231 cells expressing the MRTF-ΔSRF, comparted to wild type or MRTF-ΔSAP demonstrated a dramatic reduction in tumor initiation ^38^. Evaluating Ki67 expression in MRTF-ΔSAP tumors that were evaluable showed a significant reduction in Ki67 cells (**Fig 7E**, p<0.05).

**Figure 7.**
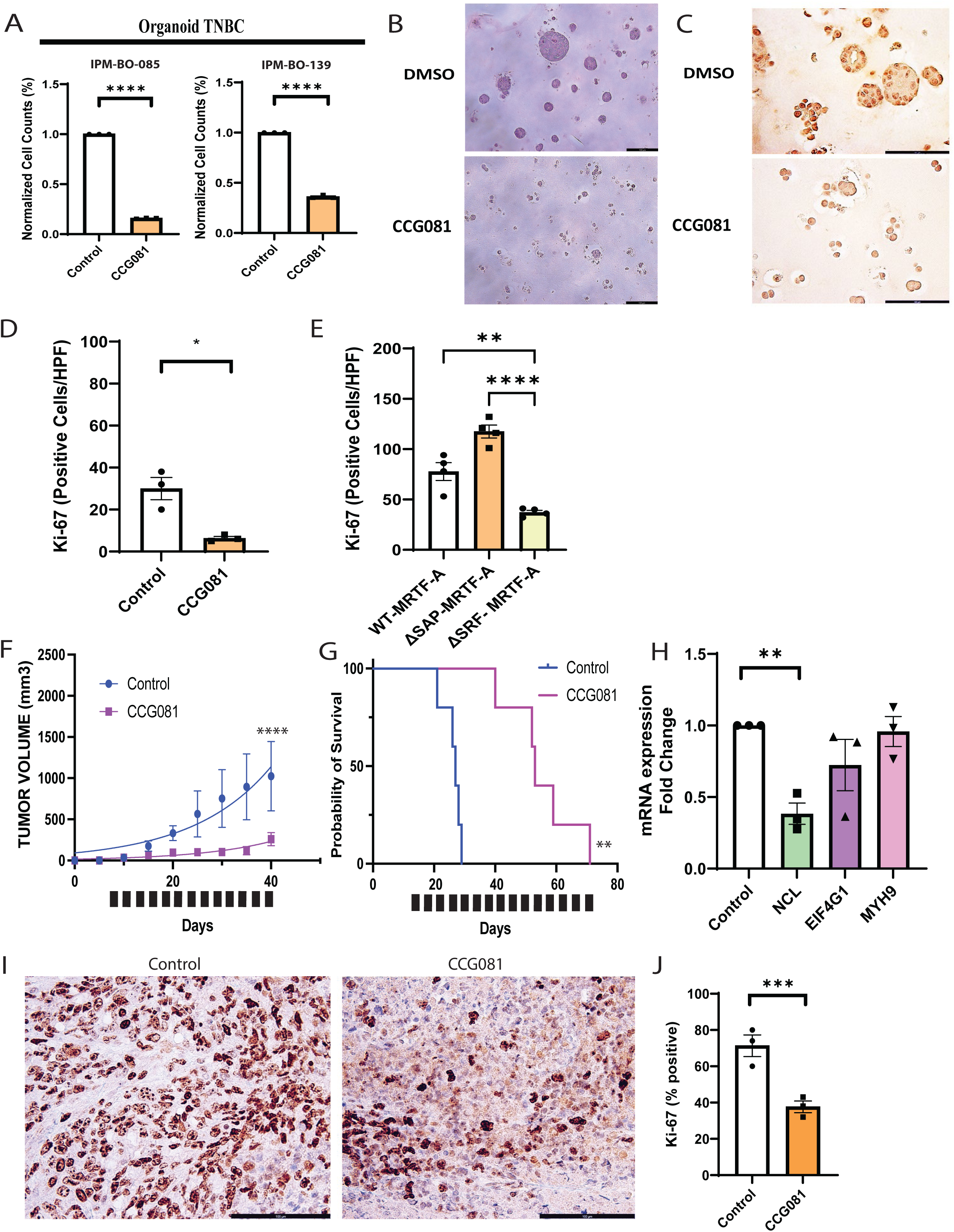
CCG257081 growth arrest in tumor models. **A-D.** Normalized cell counts, hematoxylin-eosin stain, and Ki67 IHC, with quantification from two triple negative breast cancer cell (TNBC) organoid cultures treated with DMSO (control) or CCG257081. **E.** Quantification of Ki67 stain from tumor MDA-231 cells expressing the indicated MRTF-A constructs. **F-G.** Tumor growth and overall survival curves for PT340 xenografts treated with 20 mg/Kg CCG257081 or vehicle (DMSO) control daily (black bars indicate treatment window). **H.** qRT-PCR-based relative mRNA expression for the indicated genes in control and CCG257081-treated tumors. **I-J.** Representative images and quantification of Ki67 staining in control and CCG257081 treated tumors, 6 to 8 independent sections of 3 independent tumors for each treatment group. *P<0.05, **P<0.01, ***P<0.001. HPF - High Power Field

To determine how CCG257081 performed in an *in vivo* setting, we treated PT340 ovarian cancer xenografts with CCG257081. CCG257081 treatment was associated with a prolonged restriction in tumor growth and a ∼3-fold increase in survival compared with controls (**Fig. 7F-G**, p<0.01). Evaluation of treated tumors showed CCG257081 treatment was associated with a strong reduction in NCL expression, and a decreasing trend in MYH9 and EIF4G1 expression (**Fig. 7H**, p<0.05). CCG257081-treated xenografts also had a significant reduction in Ki67 (∼ 4-fold) compared to control (**Fig. 7I-J**, p<0.01).

## Discussion

In this study, we implemented a single cell microfluidic approach to characterize the expression profile of qOvCa cells. Supporting the validity of the results, the RNA signature of qOvCa cells mirrors that seen in normal adult quiescent cells such as hematopoietic stem cells and neuronal stem cells ^33^. Like adult quiescent stem cells, qOvCa cells demonstrated significant changes that would impact protein homeostasis, including a downregulation of pathways related to mRNA processing/splicing, translation initiation, and protein/nitrogen metabolism. Interestingly, we also observed that many genes—many related to cellular metabolism—were upregulated in qOvCa. This suggests that the state of quiescence is not simply an arrested or hibernating state, but a distinct state with altered activities.

### MYH9 and nucleolin as key quiescent genes

MYH9 (myosin, heavy chain 9) is expressed in a wide range of tissues and is a member of the Myosin II motor protein super family, where it has normal physiological roles in cell migration/adhesion, cytokinesis, signal transduction and maintenance of cell structure. MYH9 has been widely reported to promote tumor progression, metastasis, and recurrence in a wide range of cancers, including gastric, colorectal, renal, esophageal, breast, and lung ^39–41^. Interestingly, MYH9 has also been shown to regulate CSC populations through activating mTOR signaling and various stem cell genes (CD44, SOX2, Nanog, CD133, and OCT4) ^42,43^.

In ovarian cancer, high expression of MYH9 in patient samples correlated with significantly worse progression-free and overall survival and more advanced FIGO staging ^44^. Corroborating our finding, a recent study demonstrated a decrease in ovarian cancer proliferation, migration, invasion, and metastasis both *in vivo* and *in vitro*, following downregulation of MYH9 with microRNA ^45^. Interestingly, MYH9 downregulation executed its antiproliferative effect through downregulation of the Wnt/β-catenin stem cell pathway and downstream cell-cycle factors ^46^, supporting our findings.

Nucleolin is a protein that is present in all growing eukaryotic cells, participating in ribosomal transcription, cell proliferation and growth. Nucleolin has been found to be implicated in multiple aspects of both DNA and RNA as well as protein metabolism ^47,48^. Nucleolin expression has been associated with tumorigenesis and angiogenesis. Nucleolin upregulation plays a critical role in molecular regulation of quiescence in hematopoiesis when it interacts with G0S2 associated proteins ^49^.

In addition to MYH9 and Nucleolin, we observed a strong induction of p27/dependency for p27 in CCG081 triggered quiescence. This is consistent for a well-established role p27 in quiescence^50^. Similarly, we observed induction of many quiescence linked factors including but not limited to NFATC4, Arora-Kinase A, and Polio like kinase ^17,51^.

### The SRF/MRTF pathway as a master regulator of quiescence

Serum is an important driver of cell proliferation, and its withdrawal can induce quiescence ^52^. This proliferative response is driven by ERK-regulated ternary complex factors (TCFs) in conjunction with SRF ^53^. Alternatively, SRF can work together with the transcription co-activator MRTF-A, regulating cytoskeleton and contractility ^54,55^. SRF/MRTF also mediates neutrophil response to LPS ^56,57^, with neutrophil activation in response to LPS linked with activation of dormant cancer cells ^58^. Indeed, increasing data also position SRF/MRTF as a critical player in cancer ^59^. Indeed, Rho kinases and the SRF/MRTF pathway are linked with cancer “stemness” and epithelial-mesenchymal transition, cell migration, and metastasis in multiple cancers ^60–63^.

Our work would suggest that, in addition to balancing proliferation and contractility, the MRTF/SRF transcription factor complex is also critically regulated in quiescent cells. As such, this axis represents an important potential therapeutic target. While transcription factors have been notoriously difficult to target therapeutically, CCG257081 successfully blocks activation of serum response element driving transcriptional activity, at least in part, by the blockade of MRTFA cytoplasmic to nuclear localization, mirroring other clinically relevant drugs such as tacrolimus, which blocks NFAT transcriptional activity by blocking cytoplasmic-to-nuclear localization ^31^.

#### Quiescence as a therapeutic target

Quiescent cells have been found to be a subset of CSC ^3–8^. CSC have long been proposed as a therapeutic target, with the goal to eradicate CSC, to overcome chemotherapy resistance ^64^. This has proven challenging to date as many critical CSC regulatory pathways, such as the Wnt pathway, are also critical in normal stem cells and thus therapeutics targeting these pathways have proven toxic. Our data suggest that an alternative treatment approach, enforcing quiescence, may be another treatment approach. Forced quiescence, while not eradicating cells and, therefore, not increasing cure, could significantly improve relapse-free survival, which could be beneficial for diseases with a rapid recurrence rate, such as ovarian or lung cancer. Importantly, our studies indicate that targeting a single component of quiescence, such as MYH9 or NCL, may be insufficient to force prolonged quiescence, as resistance develops relatively rapidly. However, targeting master regulators, such as SRF/MRTF, can have a much more profound effect. We speculate the upregulation of factors such as PLK1 and AURKA may be resistance mechanisms to drive cells back into the cell cycles. As such combination CCG081 and or PLK1 inhibitor and/or AURKA inhibitor may add additional benegit. Future studies to identify an Achilles’ heel of these cells to eradicate these quiescent state could then lead to increased cure.

### Conclusion

We have characterized qOvCa cells and identified the SRF/MRTF pathway as a critical regulator of proliferation vs. quiescence. As such, inhibition of SRF/MRTF signaling appears to drive a quiescent state in multiple cancer types. Our data implicate the SRF/MRTF as an important therapeutic target to force a quiescent cancer cell state to restrict recurrences.

## Materials and Methods

### Cell Culture

OVSAHO cells were donated by Dr. Deborah Marsh of the University of Sydney. PT412, provided by Dr. Geeta Mehta, was derived from an abdominal metastasis from a patient with platinum-sensitive high-grade serous ovarian cancer, as previously described ^65^. PT340 was used in our previous paper, with mutations as described ^21^. All cell culture media were supplemented with 10% FBS and 1% penicillin/streptomycin and cultured at 37° C and 5% CO_2_. All cell lines were tested bimonthly for the presence of mycoplasma. CCG257081 was synthesized and generously provided by Vahlteich Medicinal Chemistry Core at the University of Michigan for use in all our experiments.

### SiRNA knockdown of genes of interest

siRNA duplicates of Universal Negative Control (MISSION sigma cat# sic001), NCL (siRNA ID# SASI_Hs01_00116210 and SASI_Hs01_00116211), MYH9 (siRNA ID# SASI_Hs01_00197338 and SASI_Hs01_00197339) and EIF4G1 (siRNA ID# SASI_Hs01_00222596 and SASI_Hs01_00222597) were transfected into cell lines with Mission Transfection Reagent (Sigma #S1452) as per manufacturer’s protocol. MRTF-A siRNA (Santa Cruz, sc-43944) and MRTF-B siRNA (Santa Cruz, sc-61074) were transfected, at 10 nM concentration, with Lipofectamine RNAiMAX (Thermo Fisher, 13778100). Transfected cells were plated, in triplicate, in 2D cell culture assay and lysates harvested at 72 h post-transfection. Experiments were done in triplicate.

### Cell counting

Cell counts were performed using the Moxi Z automated counting system (Orflo Technologies) and Cassettes Type S. In cases where cell numbers were below the accuracy threshold for Moxi Z, manual hemocytometer counts were performed using trypan blue.

### Cell Cycle analysis

Cell cycle analysis was performed on OVSAHO cells lines, and the primary patient samples PT340 and PT412. Cells were fixed in 70% ethanol and incubated at −20°C for 20 minutes before being treated with RNAse A and propidium iodide (PI) for 20 minutes and run on the CytoFlex flow cytometer (Beckman Coulter), and at least 10,000 events were recorded. Cell cycle peaks were analyzed using FlowJo v10.6.2.

### Annexin V/PI staining

For apoptosis detection, cells were stained with the Annexin-V FITC apoptosis kit (BD Biosciences) according to the manufacturer’s instructions, and at least 10,000 events were analyzed on the CytoFlex flow cytometer (Beckman Coulter). The percentages of annexin V^+^, PI^+^, annexin V^+^/PI^+^, and annexin V^-^/PI^-^ cells were quantified.

### Chromatin staining assay and analysis

PT340 cells were seeded in a coverslip, then treated with either CCG257081 or cells that were previously sorted as bright/dim cells through the use of CellTrace Violet, as described in our previous protocol ^21^. They were then washed twice with phosphate-buffered saline (PBS) and fixed for 10 min at room temperature with 4% paraformaldehyde in PBS. After three washes with PBS, the cells were permeabilized for 10 min at room temperature with 0.2% Triton X-100 in PBS. Samples were then stained with Hoechst 33342 (Thermo Fisher) and mounted with ibidi mounting medium (ibidi; cat. #50001; Germany), as described in previous protocols. Following work by Parreira et al.^66^, cells were segmented and tracked over time. To evaluate the texture of cells, invariant homogeneity was calculated using the gray-level co-occurrence matrix, through the scikit-image library. Higher invariant homogeneity corresponds to a more homogeneous texture in cells, in any pixel direction.

### Fucci cell cycle reporters

HEY1 cells expressing the p27-mVenus and CDT1-mCherry FUCCI cell cycle reporter constructs were generated as previously described^34^. Cells were processed using a CytoFlex flow cytometer (Beckman Coulter), and at least 10,000 events were recorded.

### Quantitative PCR

RNA was extracted using RNeasy Mini Kit (Qiagen), and cDNA was made using SuperScript III Reverse Transcription Kit (Thermo Fisher). qPCR was performed with SYBR Green PCR Master Mix (Applied Biosystems), using standard cycling conditions. The primers used for this study are available in supplemental material.

### Microfluidics cell capture

CD133+/ALDH+ cells were isolated using FACS before 10,000 cells were loaded onto single-cell-capture microfluidic devices, using gravity flow. Microfluidic devices were imaged at Day 0 and every 24 h for 5 days. On Day 5, cells were stained with a viability stain, and single cells, whether they had divided or not, were collected via laser detachment retrieval. Single cells were snap frozen in Eppendorf tubes containing RT master mix and shipped to NYU genomic core for single-cell sequencing.

### Western Blotting

Western blotting was performed as described ^67^. Briefly, protein was lysed in RIPA buffer containing Halt Protease Inhibitor Cocktail (Thermo Fisher), sonicated, and quantified using Pierce BCA Protein Assay Kit. Thirty μg of protein was run on a 4–12% NuPAGE SDS gel (Thermo Fisher) and transferred to a PVDF membrane (Thermo Fisher). Membranes were incubated overnight with 1:1000 anti-CDKN1B (Abcam, ab32034), or 1:5000 anti-GAPDH (Proteintech cat# 60004-1-1g) antibodies in 5% skim milk. Membranes were washed in TBST, then incubated for 1h with 1:10,000 anti-mouse HRP or anti-rabbit HRP (Cell Signaling) and rewashed with TBST. Visualization was performed with ECL Plus Western Blotting Substrate (Pierce). Densitometry and quantification were subsequently performed with ImageJ.

### In vivo model

Six-week-old female NSG (NOD.Cg-Prkdc^scid^) mice were acquired from the Jackson Laboratory and allowed to acclimate 1 week in the animal facility before any intervention was initiated. All experimental procedures were conducted with the guidelines set by the Institute for Laboratory Animal Research of the National Academy of Sciences. For our oridonin and blebbistatin *in vivo* experiment, 300,000 PT340 cells were xenografted into the subcutaneous space of mice and grown until reaching approximately 100 mm^3^, at which point, the mice were treated with DMSO and either 20 mg/kg oridonin or 2 mg/kg blebbistatin daily. The mice were euthanized, and their tumors were paraffin embedded and processed for IHC. For the CCG257081 treatments, one was done to retrieve tumors from where the 300,000 PT340 cells had been injected subcutaneously, and the same parameters of tumor retrieval were used as in the previous experiments. One week after tumor injection, the animals were treated with CCG257081 at a dose of 20 mg/kg daily for 3 weeks. Mouse weight was followed once a week, and tumor volume was measured twice weekly. For the second experiment, to assess survival, 300,000 PT340 cells were injected intraperitoneally (IP). One week after tumor injection, the animals were treated with CCG257081 at a dose of 20 mg/kg daily for 3 weeks. Mouse weight was followed once a week, and the criteria for euthanasia were as previously published ^21,68^. Briefly, these were: (i) an increase of 1 cm of abdominal perimeter and/or (ii) changes in physiology and behavior (body weight, external pH or physical appearance); (iii) lower response to stimulation (inability to reach food and water, lethargy or decreased mental awareness, labored breathing, or inability to remain upright).

### Immunohistochemistry

Fresh tumors were harvested in linear growth phase (1,000 mm^3^) and embedded in formalin. Then, they were processed and embedded in paraffin 5 mm in thickness, as in previously described protocols ^21,69^. Primary anti-rabbit Ki67 (1:500, Abcam #ab15580) was incubated overnight at 4° C. Subsequently, slides were incubated with a ready-to-use peroxidase-labeled anti-mouse HRP or anti-rabbit AP (Cell Signaling Technology). Signal was visualized with DAB staining solution according to the manufacturer’s instructions and counterstained using hematoxylin. Ki67 was performed on six to eight independent sections of three independent PT340 cell-derived tumors. Images were captured on an Olympus BX41 fluorescent microscope with a 12 MB digital camera at 16-bit depth/300 dpi. Total stain area/low power field (100), was defined by pixel area (X:Y 1:1).

### Immunofluorescence

For proliferation detection by EdU staining, cells were cultured with 10 µM EdU for 24 hours and then fixed with 2% paraformaldehyde at 4°C for 30 minutes. Staining was performed using the Click-iT PLUS EdU Alexa Fluor 488 Imaging Kit (Life Technologies) and detected according to the manufacturer’s protocol. Coverslips were mounted onto slides and imaged using a Nikon A1 microscope, as previously reported ^70^.

### Statistical analysis

Statistically analysis was conducted using GraphPad Prism (10.1.0). For scRNAseq analysis, MNNCorrect and ComBat were used to confirm the absence of a batch effect. Afterward, paired t-tests were performed to compare the quiescent vs. non-quiescent genes. Data were analyzed through two-tailed Student’s t-tests or one-way ANOVA. A p value of less than 0.05 was considered to be statistically significant. For *in vitro* studies (cell counts, viability, siRNA knockdown) a minimum of three replicate experiments (n ≥ 3) were used for statistical analysis.

## Data availability statement

The data analyzed in this study were obtained from Gene Expression Omnibus. RNA Sequencing data are processed by GEO, Submission entry MvWV4Omg. Other data generated in this study are available upon request from the corresponding author.

## Acknowledgments

This work was supported by NIHR01CA278100 and NIHR01CA203810. The authors would like to thank the Institute for Precision Medicine, a partnership of the University of Pittsburgh and UPMC, for providing the breast cancer patient derived organoids used in these studies.

## Authors’ Contributions

S. Panesso-Gomez: Conceptualization, data curation, formal analysis, investigation, methodology, writing–original draft. A.J. Cole: Data curation, formal analysis, supervision, investigation, writing–original draft. A. Wield: Formal analysis, methodology. V.I. Anyaeche: Investigation. J.S. Shah: Formal analysis, methodology. Q. Jiang: Data curation, investigation. T. Ebai: Formal analysis, investigation. A.C Sharrow: Investigation. G. Tseng: Investigation. E. Yoon: Investigation. D. D. Brown: Investigation. A.M. Clark: Investigation. S.D. Larsen: Resources. D. Gau: Resources, investigation. P. Roy: Resources. KN Dahl: Resources, investigation. L Tram: Investigation H. Jiang: Resources P.F. McAuliffe: Resources A.V. Lee: Resources. R.J. Buckanovich: Conceptualization, resources, formal analysis, supervision, funding acquisition.

---

**Supplemental Table 1.**
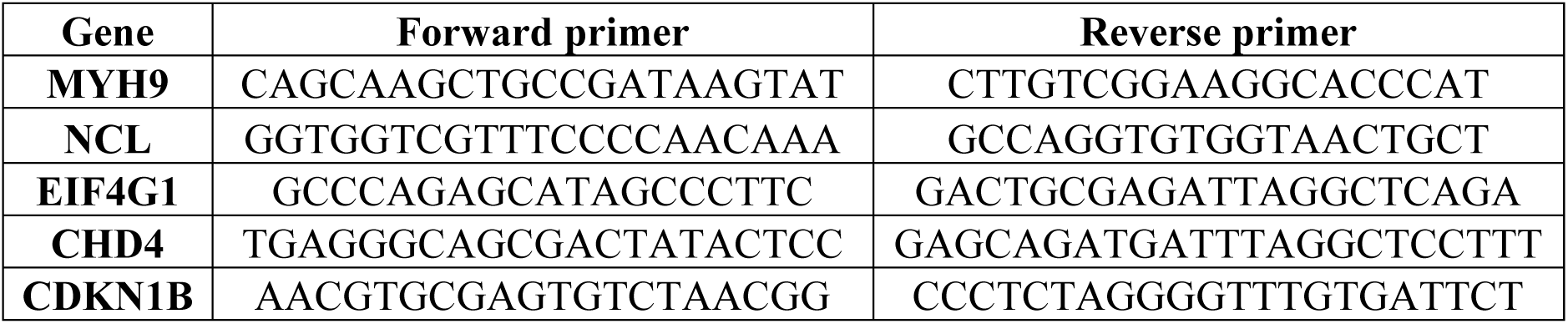
Primers used in this study.

**Supplemental Figure 1.**
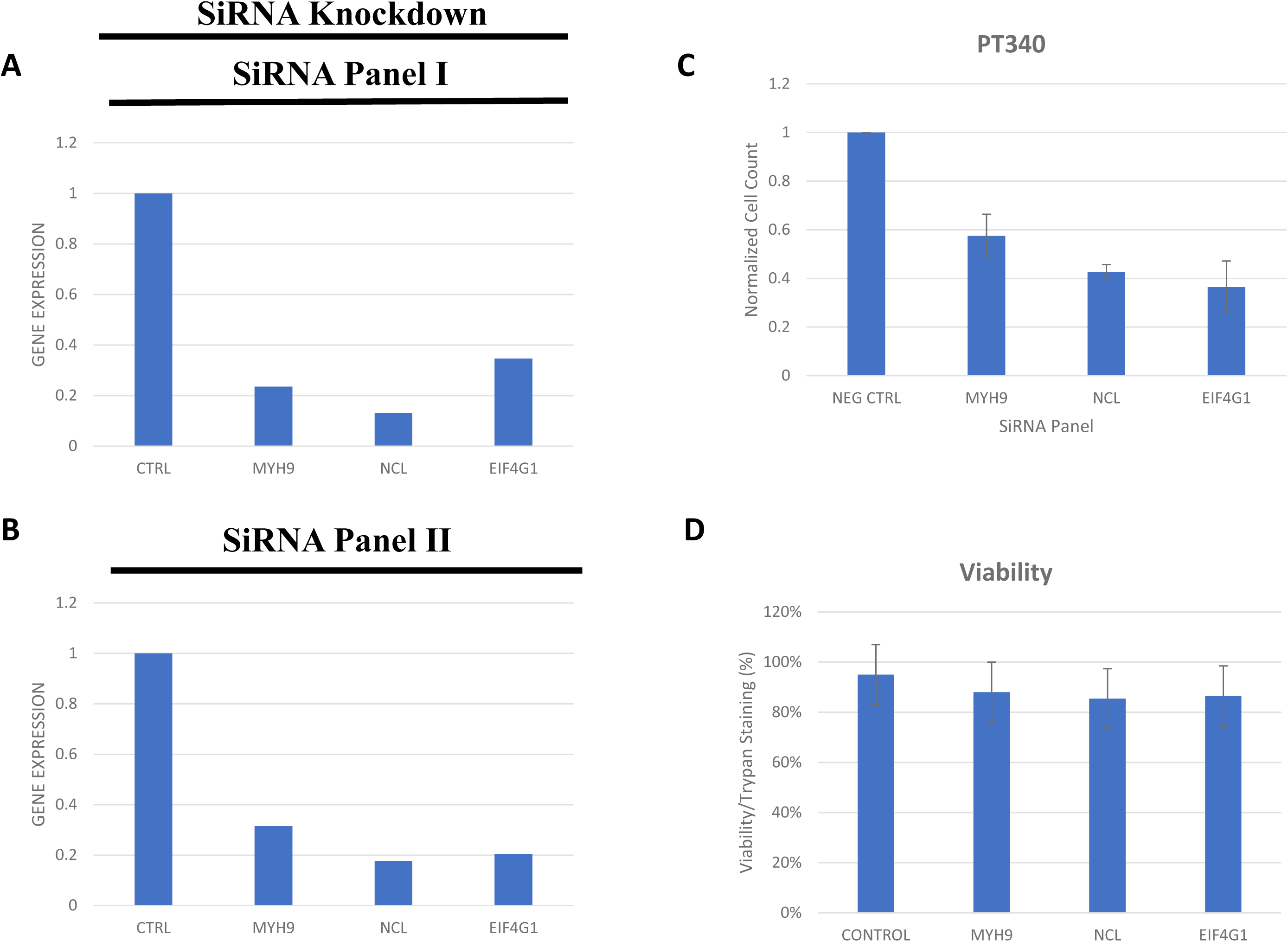
Validation of SiRNA knockdown of Microfluidics Chip Genes. **A-B.** Relative mRNA expression of the indicated of genes of interest assessed qRT-PCR. **C.** Normalized cell counts of in PT340 cell line with the indicated SiRNA knockdowns. **D.** Viability percentage in PT340 cell line with the indicated SiRNA knockdowns. Control is scrambled siRNA.

**Supplemental Figure 2.**
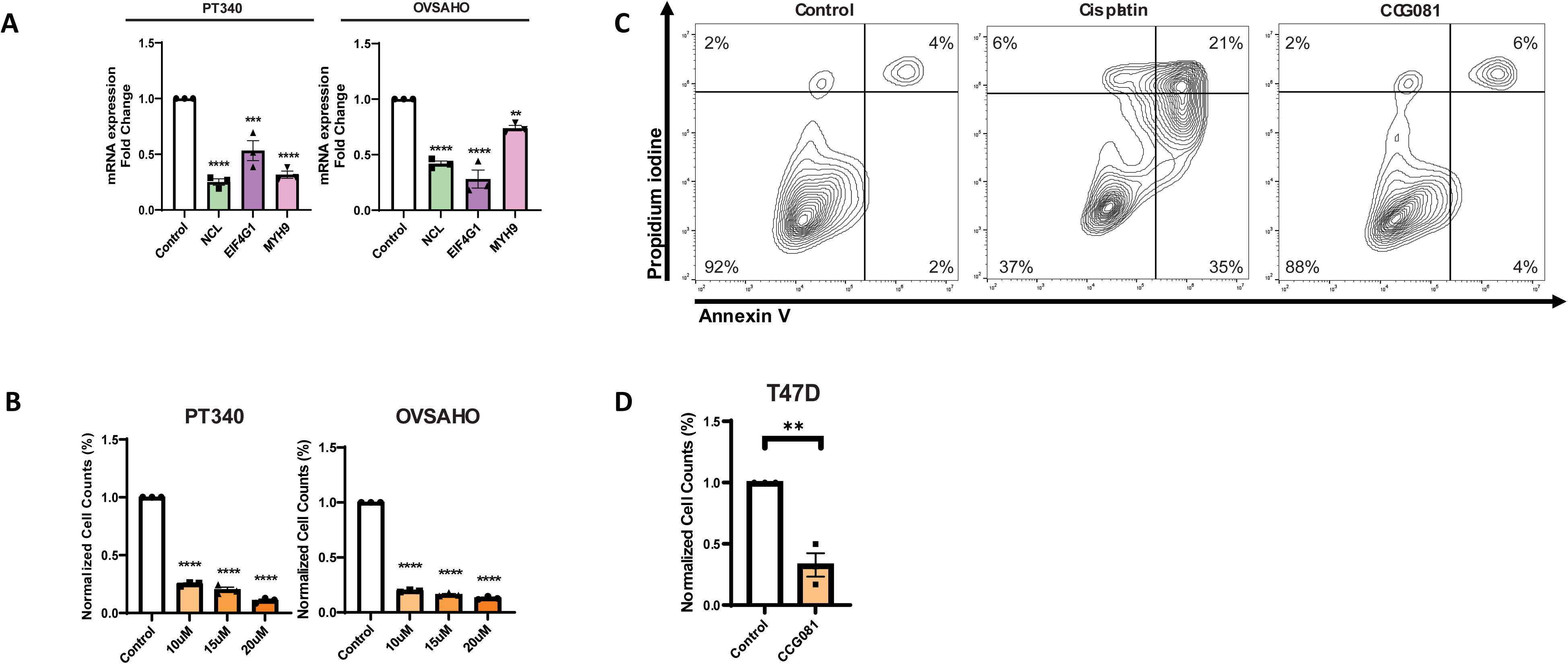
Validation of CCG257081 treatment. **A.** Relative mRNA expression of the indicated genes in control and CCG257081 treated PT340 cells. **B.** Normalized cell counts of PT340 and OVSAHO following CCG257081 treatment. **C.** Viability using Annexin V-PI staining by flow cytometry after 15uM CCG27081 or 3uM cisplatin treatment. **D.** Normalized ell count of control and CCG257081 treated breast cancer cell line T47D **p<0.01, ***p<0.001, ****p<0.0001.

**Supplemental Figure 3.**
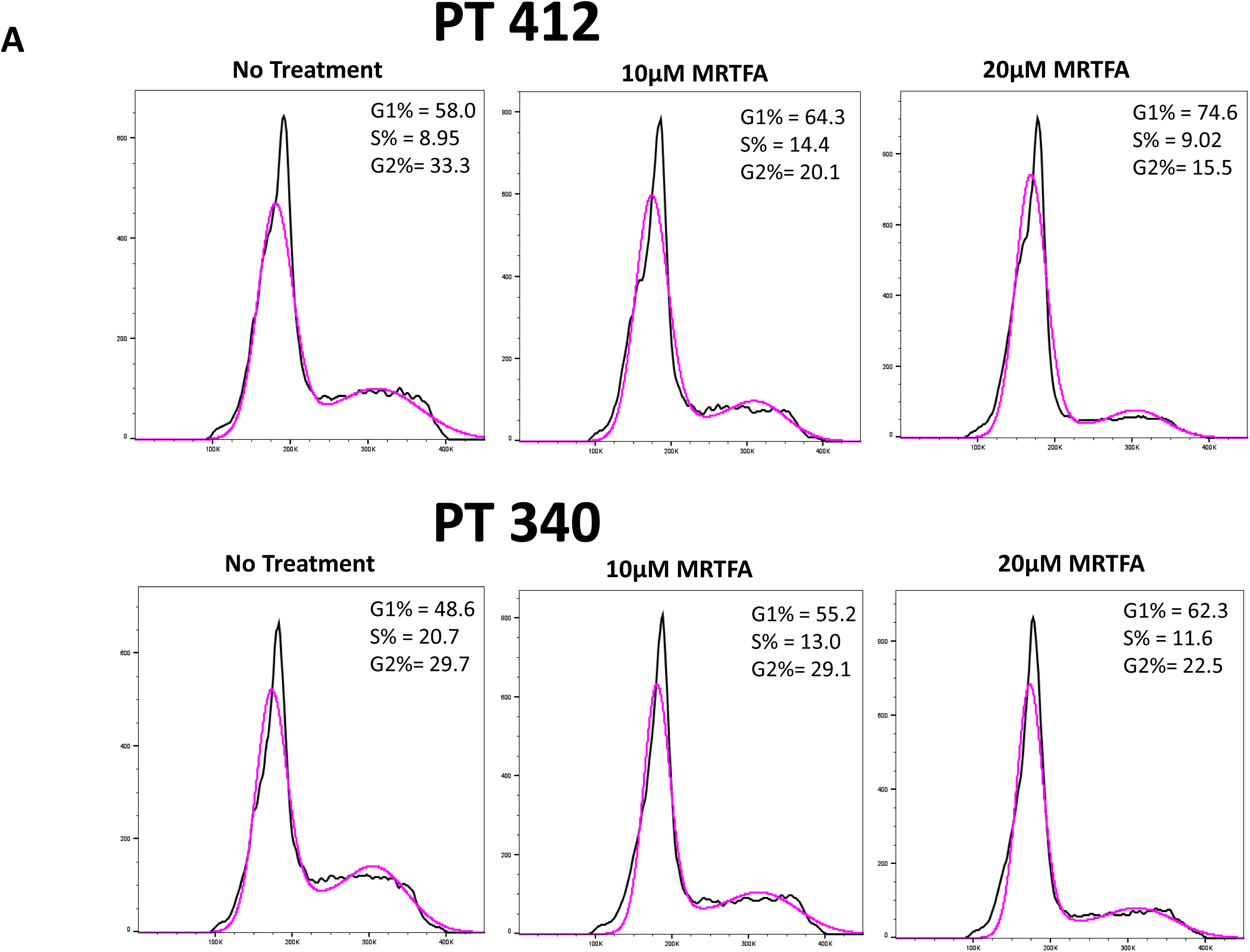
Impact of CCG257081 treatment on Cell Cycle. **A.** PI cell cycle analysis in PT412 and PT340 cells with CCG257081 treatment at a dose curve (10uM and 20uM) in two different ovarian cancer cell lines PT340 and PT412. Black line is raw counts and pink line is software normalized curves.

**Supplemental Figure 4.**
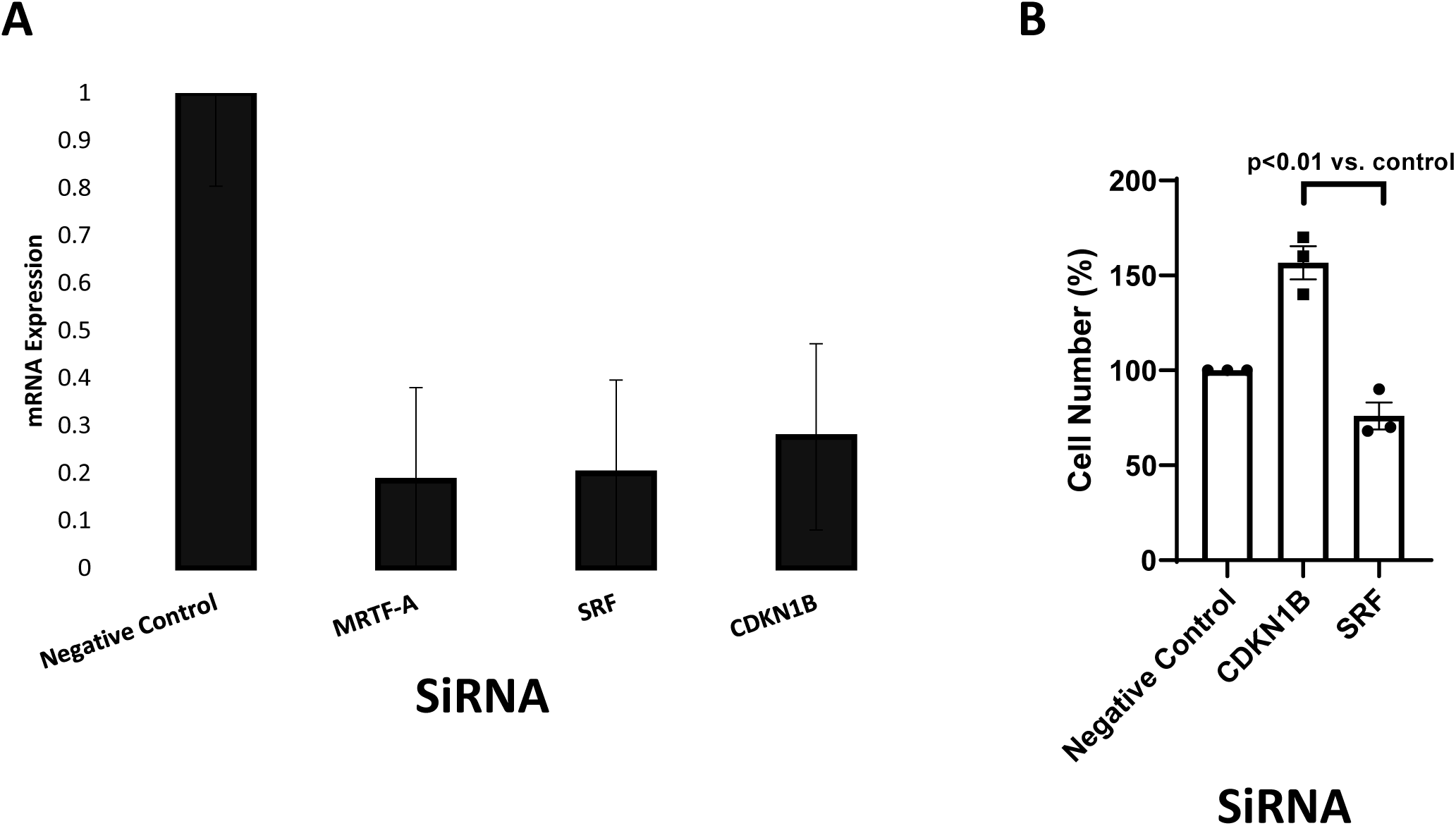
Validation of SiRNA knockdown of different factors in the MRTF/SRF pathway. **A.** qRT-PCR assessed relative mRNA expression levels with the indicated SiRNA. **B.** Relative cell number following SiRNA knockdown of p27(CDKN1B) and SRF compared to scramble control.

**Supplemental Figure 5.**
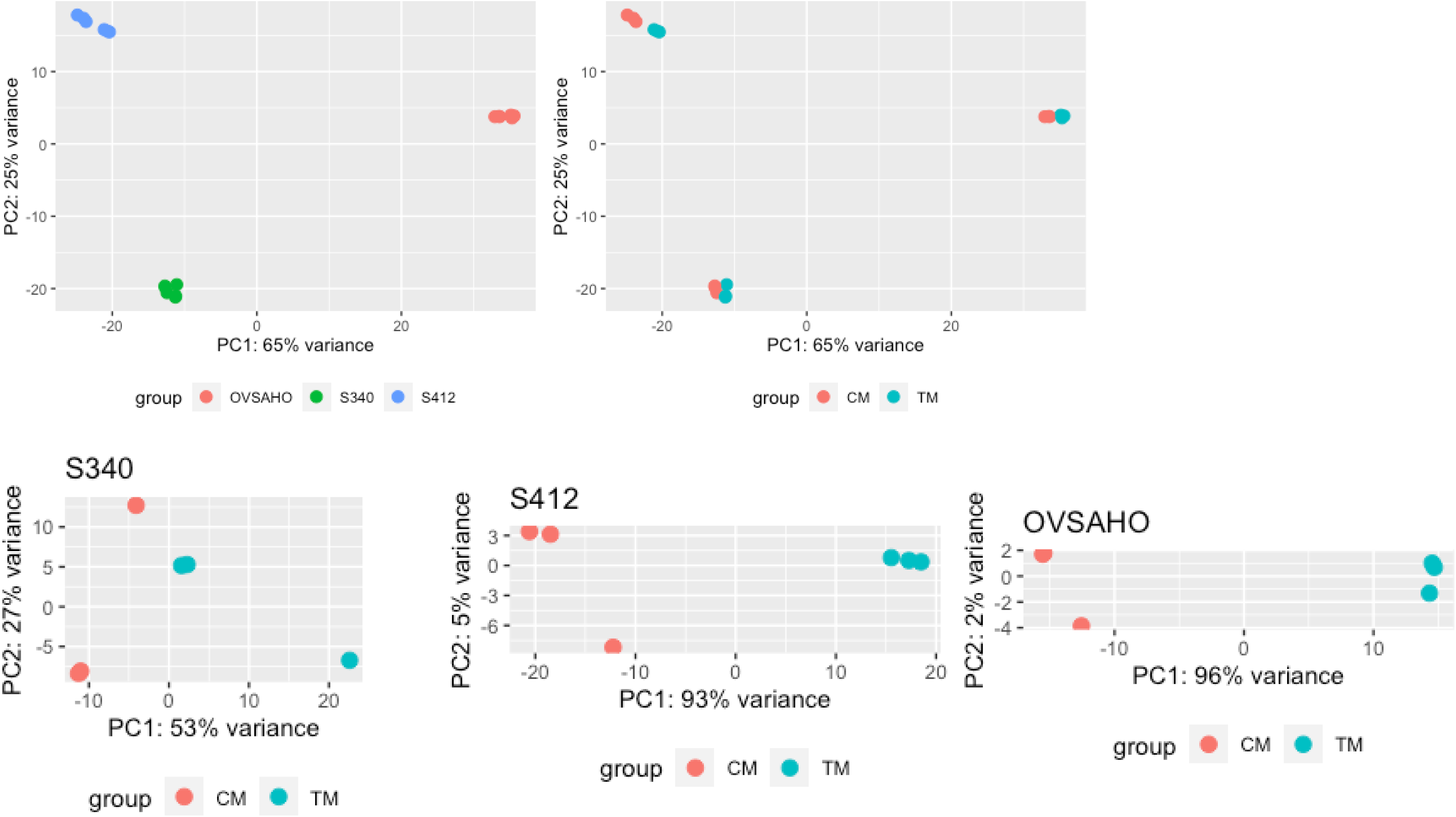
Validation of RNASeq of CCG257081 treated cells. **A.** Principal component analysis (PCA) of PT340, PT412 and OVSAHO RNA-seq datasets.

**Supplemental Figure 6.**
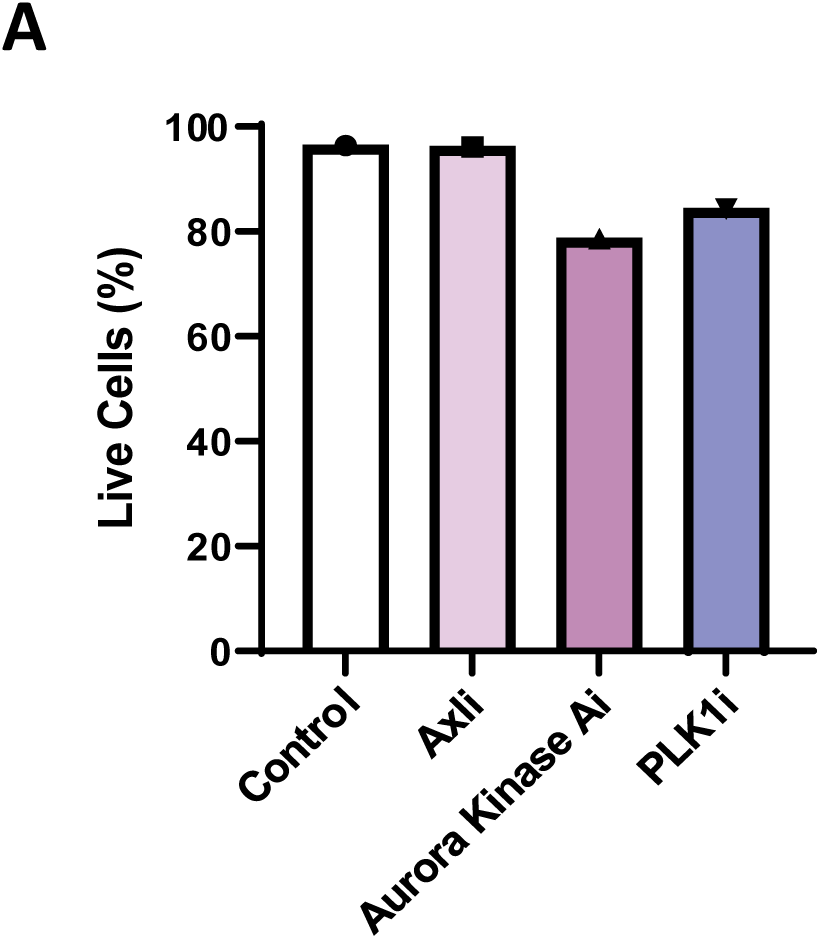
Use of inhibitors of RNAseq targets. **A.** Viability of cells treated cells with either AXL (Bemcetinib), Aurora Kinase-A (AURKA, Alisertib) or Polio-lie kinase 1 (PLK1, Onvasertib) inhibitors.

**Supplementary Table 1.**
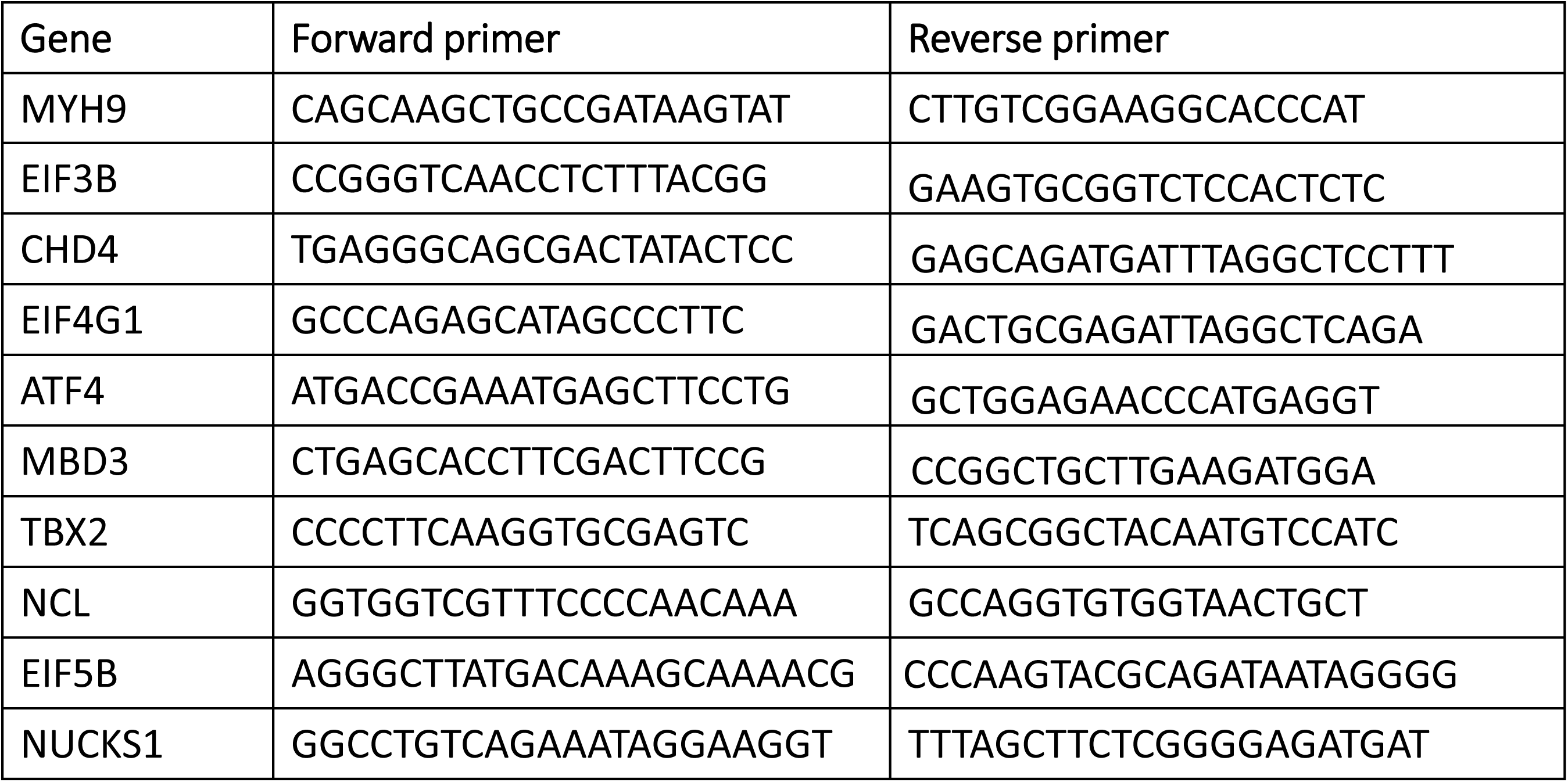
PCR Primers Used.

